# An efficient and robust tool for colocalisation: Pair-wise Conditional and Colocalisation (PWCoCo)

**DOI:** 10.1101/2022.08.08.503158

**Authors:** Jamie W Robinson, Gibran Hemani, Mahsa Sheikhali Babaei, Yunfeng Huang, Denis A Baird, Ellen A Tsai, Chia-Yen Chen, Tom R Gaunt, Jie Zheng

## Abstract

Genetic colocalisation is an important tool to test for shared genetic aetiology and is commonly used to strengthen causal inference in genetic studies of molecular traits and drug targets. However, the single causal variant assumption of the original colocalization method is a considerable limitation in genomic regions with multiple causal effects.

We integrated conditional analyses (GCTA-COJO) and colocalisation analyses (coloc), into a novel analysis tool called Pair-Wise Conditional Colocalization (PWCoCo). PWCoCo performs conditional analyses to identify independent signals for the two tested traits in a genomic region and then conducts colocalisation of each pair of conditionally independent signals for the two traits using summary-level data. This allows for the stringent single-variant assumption to hold for each pair of colocalisation analysis.

We found that the computational efficiency of PWCoCo is on average better than colocalisation with Sum of Single Effects Regression using Summary Stats (SuSiE-RSS), with greater gains in efficiency for high-throughput analysis. In a case study using GWAS data for multiple sclerosis and brain cortex-derived eQTLs (MetaBrain), we recapitulated all previously identified genes, which showcased the robustness of the method. We further found colocalisation evidence for secondary signals in nine additional loci, which was not identifiable in conventional GWAS and/or colocalisation.

PWCoCo offers key improvements over existing methods, including: (1) robust colocalisation when the single variant assumption is violated; (2) independent colocalisation of secondary signals, which enables identification of novel disease-causing variants; (3) an easy-to-use and computationally efficient tool to test for colocalisation of high-dimensional omics data.

## Introduction

Genome-wide association studies (GWAS) have identified thousands of complex traits associated genetic variants that have improved our understanding of disease aetiology, risk and treatment strategies ^1-3^. However, despite the large number of GWAS-derived associations for many complex traits, translational benefits have been limited ^4^. Colocalisation is a statistical technique that can leverage GWAS summary statistics to identify shared causal variants between two traits, which is often considered in conjunction with results from Mendelian randomisation (MR) as evidence of a causal relationship between two ^5-8^. In particular, the approach has proven valuable in validating MR of drug targets (gene expression or protein levels) used for target prioritisation and identification ^9-12^, where linkage disequilibrium (LD) in the region of the target can confound causal inference.

Given the importance of colocalisation, many packages and tools have been developed in this area, for example, coloc ^13^, eCAVIAR ^14^ and HEIDI ^15^. Each method has its own strengths, but a common limitation is the so-called “single variant assumption”, whereby it is assumed that each trait is associated with at most only one causal variant in the targeted genomic region. For genomic regions with a complex LD structure, the single variant assumption can lead to an increase in type II errors due to the acceptance of one of the models which does not show evidence of colocalisation. This has been acknowledged by the original authors of the coloc method, who recently integrated the Sum of Single Effects Regression using Summary Stats (SuSiE-RSS) framework to allow coloc to consider multiple causal variants within the same region ^16^. Other methods also exist for multiple causal variants ^8^; however, due to their complexity, these methods tend to be computationally intensive, require an estimate or assumption of how many signals may exist within a region, and can be inefficient when analysing large datasets. For example, eCAVIAR allows for the user to pre-specify how many distinct causal variants they think are present in a region; however, potential model misspecification can lead to false positives and cherry-picking of results.

In a previous systematic MR of plasma proteins, we devised a novel pipeline which we described as PairWise Conditional and Colocalisation (PWCoCo) analysis ^9^. This framework integrates approximate conditional analyses (as implemented in GCTA-COJO ^17^), which systematically conditions on each of the association signals within a genomic region, and applies pairwise Bayesian colocalisation analyses for each pair of independent signals (as implemented in the coloc R package ^13^). This conditional, pairwise approach allows the single variant assumption of the standard colocalisation method to hold for most cases (**Figure 1**). In that study, the PWCoCo pipeline was formed of a main script which ran coloc and GCTA-COJO as separate tools. Now, we have built a standalone tool in C++ which closely integrates coloc and GCTA-COJO – without altering the underlying mathematical formulation of these methods – to increase efficiency, usability and stability of running such analyses.

**Figure 1.**
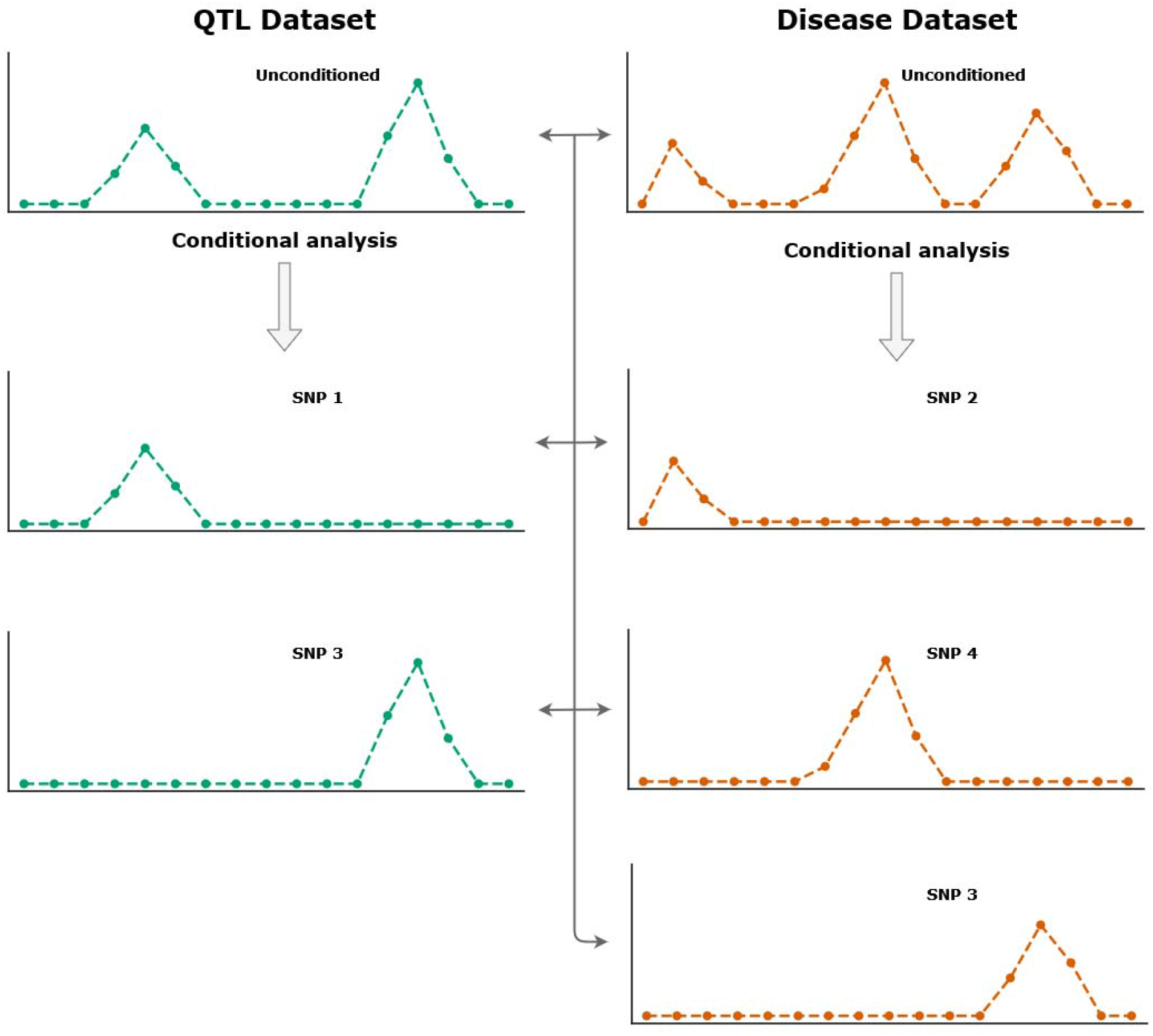
Schematic of how PWCoCo conducts conditional and colocalisation analyses in a pairwise fashion. Consider two datasets, one QTL dataset with two signals and a disease dataset with three signals. PWCoCo will conduct conditional analyses on these datasets to produce two and three new datasets, respectively, consisting of each conditionally independent signal within the region. Then, colocalisation is tested for between each of these datasets (denoted by the arrowhead lines). Therefore, in this example, a total of 12 colocalisation analyses (three QTL datasets and four disease datasets) will be run, as the unconditioned, marginal statistics are also used.

## Methods

The original PWCoCo implementation used in our systematic MR of plasma proteins consisted of an R script that made use of the coloc R package and invoked system commands to run GCTA-COJO. The major bottleneck of this approach was due to file I/O caused by running GCTA-COJO once to identify the conditionally independent signals of the target locus and then again each time to isolate each of those signals. This required the reference data to be read into memory and cleaned for each analysis as each run of GCTA-COJO was independent. Therefore, we sought to improve this by porting the colocalisation package from R to C++ and more closely integrating the conditional and colocalisation components to improve efficiency.

Here, we provide a brief overview of the PWCoCo methodology. In the first step, PWCoCo identifies all independent signals within a genomic region using a stepwise regression algorithm implemented in GCTA-COCJO: the algorithm starts by selecting the SNP with the lowest P value in the test region, then conditions all SNPs in the region on that SNP using a joint SNP model, and selects the secondary SNP with the lowest conditional P value in the joint model. This continues until the P value of the selected SNP no longer reaches the threshold defined by the user (with a default threshold of P < 5x10^-8^). As colocalisation aims to identify shared causal variants across two traits, the stepwise selection process will be conducted on the two test traits separately, to identify two sets of independent signals for the two traits in the pre-specified genomic region (e.g. *n* signals for trait 1, *m* signals for trait 2; Figure 1). For each of the constructed conditionally independent signals, colocalisation analyses are run in a pairwise manner, starting from the marginal statistics of the two traits and continuing until each pair of independent signals has been tested. Therefore, suppose the first dataset has *n* > 1 independent signals and the second dataset has *m* > 1 independent signals, PWCoCo will conduct *n* + *m* conditional analyses and (*n* + 1) × (*m* + 1) colocalisation analyses.

For input, PWCoCo accepts two summary-level GWAS datasets and a LD reference panel in PLINK^18^ 1.x format (i.e. .bed, .bim and .fam files). The LD reference panel is first used to clean the data, which consists of checking allelic information, including frequencies, against the GWAS data and second, to derive the LD structure of the given SNPs. There is no inherent restriction on which populations the data have been derived from, so long as the two GWAS datasets and the reference data are derived from the same population.

To demonstrate the efficiency of PWCoCo, we compared how our tool performed against coloc with SuSiE-RSS. We simulated summary-level data in R using European samples from the 1000 Genomes (1KG) reference panel ^19^ to generate LD-aware effects. Regions of 1Mb were randomly generated with a sample size of 10,000, between one and three distinct causal variants and between one and three shared causal variants, and the variance explained by the SNPs was set at 0.2, 0.5 or 0.8. These data were simulated using the simulateGP R package (https://github.com/explodecomputer/simulateGP) and **Supplemental Methods** contains in-depth details about how the comparison tests were conducted.

Finally, as a case study, we systematically applied PWCoCo to find potentially novel signals which colocalise between the cortex expression quantitative trait loci (eQTLs) from the MetaBrain study (n=6,601 individuals of European ancestry) ^20^ and multiple sclerosis (MS), using a GWAS from the International Multiple Sclerosis Genetics Consortium (cases = 47,429, controls = 68,374) ^21^. For each eQTL, the *cis* region around that eQTL was extracted, defined as a 1Mb window around the gene coding region. The same 1Mb windows were also extracted from the MS GWAS ^21^. We excluded those genes in the major histocompatibility complex (HMC) due to the complex LD structure in this region. Genotype data from mothers in the Avon Longitudinal Study of Parents and Children (ALSPAC) study (n = 7,927) ^22; 23^ were used as the LD reference panel data for these analyses. We also used these data to estimate the allele frequencies for the SNPs reported in the MS GWAS, which did not include this information. Any SNPs which did not contain allele frequencies after linkage were dropped from the analysis. We cross referenced our results with those generated by Baird, *et al*. ^12^, who used conventional colocalisation without fine-mapping or conditional analyses to analyse both of the same datasets. We also attempted to repeat this analysis using coloc with SuSiE-RSS.

## Results

We observed that PWCoCo runs substantially faster than the first iteration of PWCoCo described in Zheng, *et al*. ^9^ (roughly 99% faster) and coloc with SuSiE-RSS (between 82-87% faster) (**Table 1**). The runtime for SuSiE-RSS is substantially reduced if the time taken to calculate the LD matrix is not taken into consideration; however, as PWCoCo generates the underlying LD of the region from the raw genotype data, a fairer comparison is made by including that time. Separately, we saw a roughly 36% decrease in time taken to perform 200 analyses when PWCoCo can be allowed to read and prepare the reference panel once as opposed to for every analysis (10 seconds per analysis vs on average 6 seconds per analysis in the 200 analyses run).

**Table 1.** Time taken to run between 1 and 200 analyses using PWCoCo presented in Zheng, *et al*. ^9^, PWCoCo presented in this paper, and coloc with SuSiE-RSS. PWCoCo was conducted with two configurations: 1) analyses were conducted separately, each reading and cleaning the reference panel and not storing the cleaned data in memory. 2) analyses were run as a batch, such that PWCoCo only required to operate on the reference data once and stored this in memory. The bottleneck for PWCoCo is file operations on large reference data, so storing this data in memory can be expensive but will save time if that is a concern. If only a few analyses need to be conducted, then streaming the reference panel will not substantially increase runtime. As more analyses are required, then loading the reference data into memory will become more attractive due to the time it saves. Coloc with SuSiE-RSS was also run with two configurations: 1) the LD matrix was not pre-calculated. Plink was used to generate the LD matrix for the SNPs in the region using the whole 1KG phase 3 reference panel. The time Plink took to generate the matrix was included in this measure. 2) the LD matrix was pre-calculated using Plink and 1KG phase 3 reference panel, such that these runtimes measure only the time taken by coloc with SuSiE-RSS to run. Finally, PWCoCo as presented in Zheng, *et al*. did not finish within the given time (seven days).

| Analyses | PWCoCo,<br>Zheng, *et al.* | PWCoCo,<br>Robinson, *et al.* |  | Coloc with SuSiE-RSS |  |
| --- | --- | --- | --- | --- | --- |
|  |  | Sequential | Memory | Inc. LD matrix | Exc. LD matrix |
| 1 | 52.3 minutes | 9.5 seconds | 9.6 seconds | 54.3 seconds | 2.4 seconds |
| 100 | 4.2 days | 15.4 minutes | 11.4 minutes | 90.5 minutes | 4.1 minutes |
| 200 | Did not finish | 30.9 minutes | 19.9 minutes | 3.0 hours | 8.2 minutes |

In the simulation tests comparing PWCoCo’s performance to coloc with SuSiE-RSS, we observed that coloc with SuSiE-RSS tended to underestimate, while PWCoCo tended to overestimate, the number of signals which colocalised between datasets (**Table 2**). We also found that coloc with SuSiE-RSS showed better performance in identifying shared causal variants (2,699 / 5,000 = 53% to 1,223 / 5,000 = 24%) than PWCoCo (1,600 / 5,000 = 32% to 879 / 5,000 = 18%). Coloc with SuSiE-RSS also showed higher accuracy in identifying the exact shared causal variant (15% to 2% vs 0.8% to 0.3% for PWCoCo). We also compared how frequently both methods identified a variant in high LD (r^2^ > 0.8) with the exact shared causal variant(s); in this case, PWCoCo performed similarly to SuSiE-RSS, likely due to COJO’s approximate nature. The rate at which both methods found either the true or high LD proxy for the shared causal variant decreased as more causal variants were added to the simulated data (**Supplemental Table 1**).

**Table 2.** Results for simulations to test performance of PWCoCo and SuSiE. Each dataset was simulated with between one and three distinct causal variants and between one and three shared causal variants. Therefore, each dataset had between two and six causal variants. The “exact” column contains number of datasets for which that method found the exact shared causal variant(s) while the “high LD” column contains the number of datasets for which that method found a variant which was in high LD (r^2^ > 0.8) with the shared causal variant(s). Also shown is the mean number of signals across all datasets which had strong evidence of colocalisation (H4 > 80%). 5,000 datasets were simulated for each configuration of shared causal variants. **Supplemental Methods** contains information on how these simulations were conducted. **Supplemental Table 1** contains granular results split over the amount of distinct and shared causals variants.

| Shared variant(s) | SuSiE |  |  | PWCoCo |  |  |
| --- | --- | --- | --- | --- | --- | --- |
|  | Exact | High LD | Mean signals with H4 > 80% | Exact | High LD | Mean signals with H4 > 80% |
| 1 | 727 | 1942 | 0.83 | 42 | 1558 | 2.14 |
| 2 | 281 | 1539 | 1.42 | 24 | 1220 | 2.72 |
| 3 | 119 | 1104 | 1.78 | 13 | 866 | 3.13 |

In the case study analysis to determine which MetaBrain eQTLs colocalise with MS, we found that we reproduced all of the results presented in Baird, *et al*. ^12^ (**Supplemental Table 2**). In total, we found strong colocalisation evidence (H4 ≥ 80%) for 76 signals and moderate colocalisation evidence (80% > H4 ≥ 60%) for a further 82 signals (**Figure 2**). Of these 158 results, 42 were derived using non-marginal statistics in the MetaBrain dataset and five in the MS dataset. Just over half (24) of those results derived using non-marginal statistics in the MetaBrain study were non-primary signals. Full results for these analyses are given in **Supplementary Table 3**. Our results contained many genes which were not analysed in the Baird, *et al*. paper because the authors assessed colocalisation evidence for eQTLs which already had strong MR evidence between the eQTL and MS. We also found strong evidence for colocalisation (H4 ≥ 80%) for nine genes that did not show strong evidence when using the marginal statistics. These were: *MARK3, FCRL1, CHCHD2, ATP1A4, NBEAL2, EIF2AK3, STARD10, FCRL3* and *RPIA*. Of these nine genes, *ATP1A4, NBEAL2* and *EIF2AK3* had strongest evidence for colocalisation using non-primary signals highlighting the importance of considering the presence of multiple signals at a locus when conducting colocalisation.

**Figure 2.**
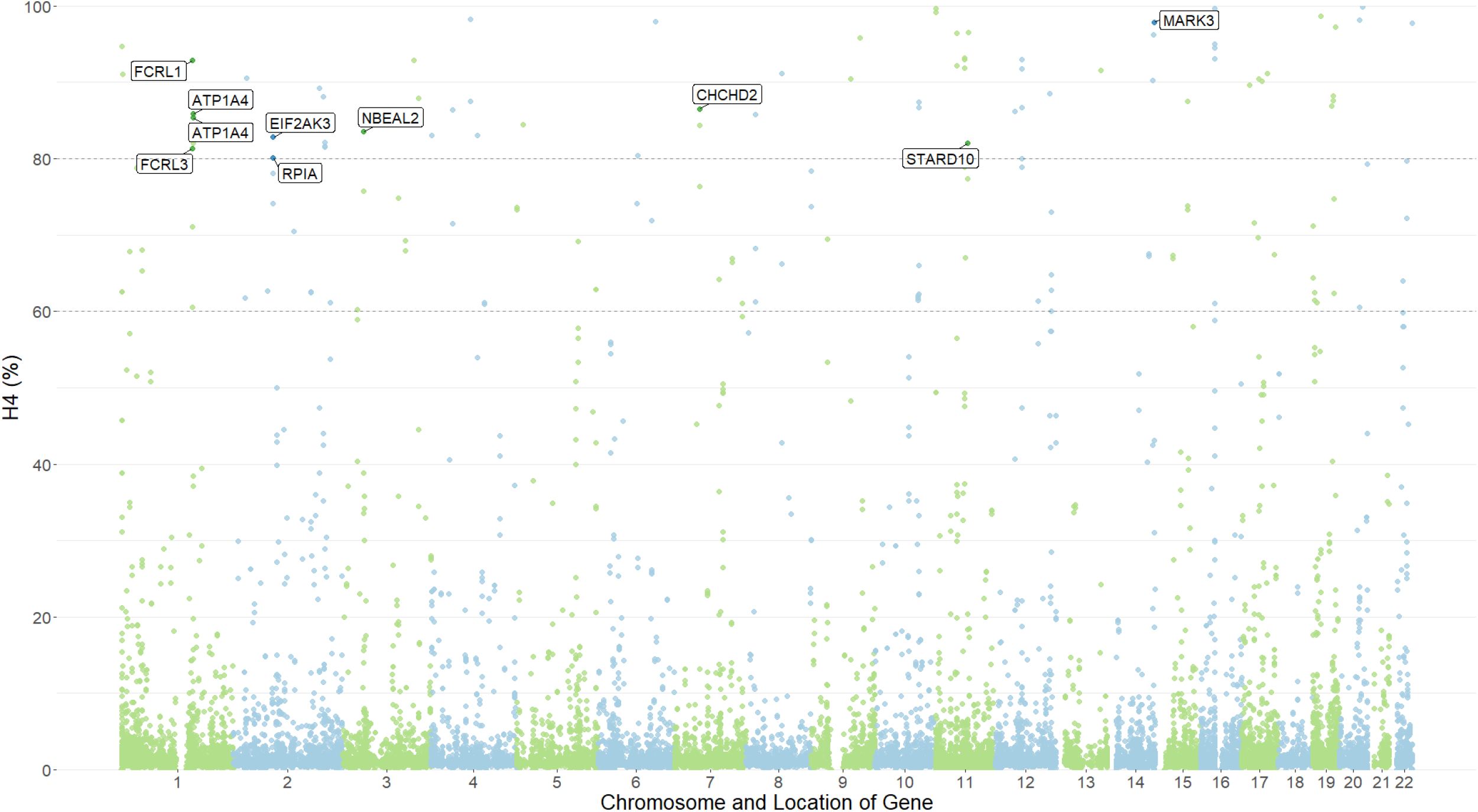
Manhattan plot of results from the systematic application of PWCoCo on cortex eQTLs and MS. Each dot represents a colocalisation analysis result. Many of the analysed loci had multiple signals in the region, either in the eQTL or MS dataset; therefore, gene names may appear more than once on the graph for secondary, tertiary, etc. signals. Only the H4 results are plotted, and genes were plotted using the start location of their gene encoding region. Labelled points are those genes with novel colocalisation.

We attempted to conduct the same analysis using coloc with SuSiE-RSS; however, we found that SuSiE-RSS failed to run due to an LD mismatch between the summary data and two reference panels (1KG phase 3 and mothers in ALSPAC). We conducted post-hoc diagnostic tests to investigate why SuSiE-RSS failed to run and found that the parameter *λ*, which is used as a measure of the inconsistency between the summary statistics and LD matrix, was high for many of the tested loci indicating large inconsistencies between the observed and estimated z-scores (smallest λ = 0.19, largest λ = 0.91, average λ = 0.59; **Supplemental Table 4**). This analysis is further detailed in **Supplemental Methods**.

## Discussion

In this paper, we have presented PWCoCo, a tool which integrates pairwise conditional analyses with Bayesian colocalisation to maintain the single variant assumption that otherwise limits such analyses. We compared how PWCoCo performed against coloc with SuSiE-RSS, a similar method to PWCoCo, using simulated data. Finally, we applied PWCoCo to ascertain which eQTLs in the MetaBrain ^20^ dataset colocalised with MS.

In the simulation tests, we observed that PWCoCo performed faster than coloc with SuSiE-RSS; SuSiE-RSS was only faster when it had access to pre-calculated LD matrices for each of the analyses. Furthermore, we made substantial improves to the efficiency of PWCoCo when compared to its first iteration presented in Zheng, *et al*.^9^.

To test for correctness, we examined the mean of how many signals had high evidence of colocalisation (H4 ≥ 80%). Each dataset was simulated with between one and three shared causal variants, and we found that while coloc with SuSiE tended to underestimate the number of signals which colocalised, PWCoCo overestimated the number of signals. Furthermore, SuSiE appeared to be the more accurate of the two methods at identifying the exact causal variant, though both methods performed similarly when allowing for finding a variant in high LD (r^2^ > 0.8) with the true causal variant. This result was to be expected given the approximate nature of COJO which PWCoCo uses and how the simulation tests slightly favoured SuSiE-RSS (discussed further below). However, it is not entirely necessary for PWCoCo to accurately tag the true causal variant given the purpose of the tool is to perform colocalisation analyses, which assumes the presence of a causal variant and is robust given a dense enough region of SNPs ^8^.

We used PWCoCo to provide evidence for colocalisation between cortex-derived eQTLs in the MetaBrain dataset ^20^ and MS ^21^ and compared our results to a previous study by Baird, *et al*. ^12^ which analysed the same datasets using only conventional colocalisation. PWCoCo found evidence of colocalisation for nine genes not found in the original analysis. Three of these genes colocalised with non-primary signals which may have gone otherwise un-examined in a typical colocalisation analysis, where some studies only use marginal statistics, despite the literature advising that this will bias results ^13; 16; 24^. Furthermore, these non-primary signals may be translationally useful, particularly in drug target identification and prioritisation, where robust genetic evidence can increase the success rate of targets in early clinical trials ^25^.

Although coloc with SuSiE-RSS performed well in our simulation study, this method is sensitive to LD mismatch in real-data analyses and thus did not provide any meaningful results when applied to the same MetaBrain and MS GWAS datasets case study. SuSiE-RSS does not require the reference data to be derived from the same sample as the summary data it is recommended as inconsistences between z-scores and the derived LD matrix can result in difficulties conducting such analyses ^26^. We showed that among the 132 genes which showed strong colocalisation evidence using PWCoCo, the lambda parameter (used to measure inconsistences between the z-scores and LD matrix) ranged from 0.19 to 0.91 in the MetaBrain dataset using either the 1KG or mothers in ALSPAC reference panels. These number were much higher than the lambda tested in the original SuSiE-RSS publication (less than 0.01), and the fine-mapping subsequently failed for 73% the loci tested; the remaining genes for which SuSiE-RSS did converge provided spurious results as evidenced by finding credible sets consisted of an implausible number of SNPs, or for which consisted of an infinite log10 Bayes factor. Furthermore, the authors of SuSiE-RSS caution against using meta-analysed datasets, where the inclusion of SNPs not measured in all of the constituent datasets might produce spurious results which is a considerable limitation of the methodology as meta-analysis is an attractive technique to increase sample sizes and thus statistical power ^20; 27-29^.

While PWCoCo benefits from the computation efficiency of GCTA-COJO for conditional analysis, there are also several limitations that comes with the implementation of GCTA-COJO. First, it is suggested that GCTA-COJO, and therefore PWCoCo, requires a minimal of 4,000 reference samples for LD calculation ^17^. This is, however, no longer a difficulty with publicly available resources such as UK10K, the haplotype reference consortium (HRC) and UK Biobank. Second, an important distinction between of PWCoCo and coloc with SuSiE-RSS is that GCTA-COJO is an approximate conditional method, while SuSiE-RSS directly performs fine-mapping on the marginal GWAS summary statistics. Therefore, SuSiE-RSS is expected to perform better at determining the true causal variant or variants at a locus. We observed this phenomenon in our simulation study; although, our simulation study is biased toward SuSiE-RSS because the data were simulated using an LD matrix derived from the 1KG phase 3 reference panel ^19^ and the exact LD matrix was then directly provided to coloc with SuSiE-RSS, whilst PWCoCo used the raw genotype data for conditional analysis. Furthermore, this reference panel has a sample size of only 2,504 which may leads under performance in finding the true causal variant for PWCoCo.

Strengths of PWCoCo include its computational speed and efficiency as evidenced by the simulation studies. PWCoCo runs single or few analyses quickly but can slow down when many analyses need to be conducted, as may be common for determining genome-wide colocalisation evidence for molecular traits. This is due to the bottleneck caused by parsing and cleaning the reference data; therefore, we developed a modified method of running PWCoCo which allows for this process to be run once, saving the user time in conducting large-scale analyses which rely on the same reference data. PWCoCo is also characterised by its flexibility, due to many user-alterable parameters (these are documented on the PWCoCo GitHub repository, https://github.com/jwr-git/pwcoco), and ease of use, requiring only summary-level data and a reference panel in PLINK 1.x binary format ^18^. Furthermore, PWCoCo is also ancestry agnostic so long as the summary data and reference panel are derived from the same population. Finally, PWCoCo is robust to the issues we observed when attempting to run SuSiE-RSS on the MetaBrain dataset, namely: mismatched z-scores and LD matrices and the requirement to remove SNPs in meta-analysed datasets which are not measured in all constituent studies.

Overall, PWCoCo is an efficient, easy-to-use tool that combines conditional and colocalisation analyses to both increase robustness of Bayesian coloc results and to allow analysis of non-primary signals. We have shown that PWCoCo performs faster than coloc with SuSiE-RSS. In a case study, PWCoCo replicated previously published results when applied to GWAS of MS and brain-derived eQTLs ^12^. Furthermore, PWCoCo found novel colocalisation evidence for additional non-primary eQTLs which went unanalysed in the original publication. Investigators should consider integrating PWCoCo into their analytical pipelines in place of colocalisation, especially when many large-scale analyses need to be conducted or when using meta-analysed datasets.

## Supporting information

Supplemental Results

## Acknowledgements

This work was carried out using the computational facilities of the *Advanced Computing Research Centre*, University of Bristol - http://www.bristol.ac.uk/acrc/.

## Competing Interests

JWR and TRG received funding from Biogen, Inc. for other projects. YH, DAB, EAT and C-YC are employees of Biogen, Inc.

## Code Availability

PWCoCo can be found on GitHub: https://github.com/jwr-git/pwcoco. Instructions for installation and operating parameters are also found on the GitHub. The simulateGP R package we used to generate the simulated datasets can also be found on GitHub: https://github.com/explodecomputer/simulateGP.

## Notes

https://github.com/jwr-git/pwcoco

