## Supplemental Methods for "An efficient and robust tool for colocalisation: Pair-wise Conditional and Colocalisation (PWCoCo)"

### Simulation Study

To conduct the simulation tests, we utilised an R package called simulateGP (<https://github.com/explodecomputer/simulateGP>) which allows for simulation linkage disequilibrium (LD)-aware summary-level genome-wide association study (GWAS) data. We used 1000 Genomes (1KG) phase 3 reference panel ^1^ with European only samples to generate the underlying LD structure of the data. Regions were split into broadly independent LD blocks as reported by Berisa and Pickrell ^2^, with three regions randomly selected from each chromosome (excluding the major histocompatibility complex (HMC)). Tests were conducted on BlueCrystal 4, a high-performance computing system at the University of Bristol, using the same resources for PWCoCo and coloc with Sum of Single Effects Regression using Summary Statistics (SuSiE-RSS): eight cores (2.4 GHz Intel E5-2680 v4 (Broadwell) CPUs) with 48 GB of RAM. Both PWCoCo and SuSiE-RSS were allowed to access as many threads as allowable (for example, PWCoCo makes use of OpenMP ^3^ to speed up certain processes).

Regions were generated with between one and three distinct causal variants and one and three shared causal variants. Therefore, each dataset had between two and six causal variants. Data were simulated with a sample size of 10,000 and the variance explained by the SNPs for the trait were set to either 0.2, 0.5 or 0.8. Causal variants were sampled to be within ±500Kb of each other. The selection of coefficient on the trait was set to zero as is default for the simulatedGP package. Finally, each dataset was generated and tested three times.

Coloc with SuSiE-RSS was run with mostly default parameters, except for a maximum limitation on the number of iterations to check for convergence, which was set to 1 million iterations. PWCoCo has a limit which if the initial colocalisation on the marginal statistics exceeds, the tool terminates early (default H4 ≥ 80%). This was disabled for the simulation tests, whereas all other parameters were the default.

### Application of SuSiE-RSS on MetaBrain eQTLs

We attempted to run coloc with SuSiE-RSS using the MetaBrain expression quantitative trait loci (eQTL) and multiple sclerosis (MS) datasets, but SuSiE-RSS failed to run when using these data. The authors of SuSiE-RSS, in their paper, discuss reasons why this might happen and how to diagnose such issues ^4^. Zou, *et al.* advise that SuSiE-RSS can fail when there are discrepancies between the summary and reference data, which primarily occurs due to a mismatch between the given z-scores and the LD matrix but can also occur due to an allele mismatch. The authors provide a function (“estimate_s_rss”) which can estimate a measure of the inconsistency between the summary statistics and the LD matrix (λ), where λ is estimated by maximum likelihood and is bound between 0 and 1 for high and low consistency, respectively. Selecting the eQTL loci which showed strong evidence of colocalisation (H4 ≥ 80%) in the PWCoCo analysis (n = 132), we generated λ using both the 1KG phase 3 and ALSPAC reference panels. We found that λ ranged between 0.19 and 0.91 (mean = 0.59) and SuSiE-RSS failed to generate a credible set for 98 loci.

For those loci which SuSiE-RSS succeeded in producing a credible set, these either consisted of an improbable 43 fine-mapped variants or consisted of credible sets whose log10 Bayes factor was infinite. Zou, *et al*. provide a potential reason for these results by advising caution if the summary statistics are derived from a meta-analysis whereby some SNPs are not analysed in all studies and thus have a lower sample size than others, as this can bias fine-mapping results ^4^. Therefore, as this applies to the MetaBrain dataset, which is a meta-analysis of 14 datasets, we dropped SNPs which did not have the maximum sample size of 6,601. However, none of the SNPs at any of the tested loci had the maximum sample size of 6,601.
